# Granatum: a graphical single-cell RNA-Seq analysis pipeline for genomics scientists

**DOI:** 10.1101/110759

**Authors:** Xun Zhu, Thomas Wolfgruber, Austin Tasato, David G. Garmire, Lana X Garmire

**Affiliations:** Graduate Program in Molecular Biology and Bioengineering, University of Hawaii at Manoa, Honolulu, HI 96816; Epidemiology Program, University of Hawaii Cancer Center, Honolulu, HI 96813; Department of Electrical Engineering, University of Hawaii at Manoa, Honolulu, HI 96816

**Keywords:** single-cell, gene expression, graphical, normalization, clustering, differential expression, pathway, pseudo-time, software

## Abstract

**Background:** Single-cell RNA sequencing (scRNA-Seq) is an increasingly popular platform to study heterogeneity at the single-cell level.

Computational methods to process scRNA-Seq have limited accessibility to bench scientists as they require significant amounts of bioinformatics skills.

**Results:** We have developed Granatum, a web-based scRNA-Seq analysis pipeline to make analysis more broadly accessible to researchers. Without a single line of programming code, users can click through the pipeline, setting parameters and visualizing results via the interactive graphical interface Granatum conveniently walks users through various steps of scRNA-Seq analysis. It has a comprehensive list of modules, including plate merging and batch-effect removal, outlier-sample removal, gene filtering, geneexpression normalization, cell clustering, differential gene expression analysis, pathway/ontology enrichment analysis, protein-network interaction visualization, and pseudo-time cell series construction.

**Conclusions:** Granatum enables broad adoption of scRNA-Seq technology by empowering the bench scientists with an easy-to-use graphical interface for scRNA-Seq data analysis. The package is freely available for research use at http://garmiregroup.org/granatum/app

## Background

Single-cell high-throughput RNA sequencing (scRNA-Seq) is providing new opportunities for researchers to identify the expression characteristics of individual cells among complex tissues. From bulk cell RNA-Seq, scRNA-Seq is a significant leap forward. In cancer, for example, scRNA-Seq allows tumorous cells to be separated from healthy cells [1], and primary cells to be differentiated from metastatic cells [2]. Single-cell expression data can also be used to describe trajectories of cell differentiation and development [3]. However, analyzing data from scRNA-Seq brings new computational challenges, e.g., accounting for inherently high drop-out or artificial loss of RNA-expression information [4,5].

Software addressing these computational challenges typically requires the ability to use a programming language like R [5,6], limiting accessibility for biologists who only have general computer skills. Existing workflows that can be used to analyze scRNA-Seq data, such as Singular (Fluidigm, Inc., South San Francisco, CA, USA), Cell Ranger (10x Genomics Inc., Pleasanton, CA, USA), and Scater [7], all require some non-graphical interactions. They also may not provide a comprehensive set of scRNA-Seq analysis methods. To fill this gap, we have developed Granatum, a fully interactive graphical scRNA-Seq analysis tool. Granatum takes its name from the Latin word for pomegranate, whose copious seeds resemble individual cells. This tool employs an easy-to-use web-browser interface for a wide range of methods suitable for scRNA-Seq analysis: removal of batch effects, removal of outlier cells, normalization of expression levels, filtering of under-informative genes, clustering of cells, identification of differentially expressed genes, identification of enriched pathways/ontologies, visualization of protein networks, and reconstruction of pseudo-time paths for cells. Our software empowers a much broader audience in research communities to study single-cell complexity by allowing the graphical exploration of single-cell expression data, both as an online web tool (from either computers or mobile devices) and as locally deployed software.

## Implementation

### Overview

The front-end and the back-end of Granatum are written in R [8] and built with the Shiny framework [9]. A load-balancer written in NodeJS handles multiple concurrent users. Users work within their own data space. To protect the privacy of users, the data submitted by one user is not visible to any other user. The front-end operates within dynamically loaded web pages arranged in a step-wise fashion. ShinyJS [10] is used to power some of the interactive components. It permits viewing on mobile devices through the reactivity of the Bootstrap framework. To allow users to redo a task, each processing step is equipped with a reset button. Bookmarking allows the saving and sharing of states.

### Interactive widgets

Layout and interactivity for the protein-protein interaction (PPI) network modules is implemented using the visNetwork package [11]. Preview of user-submitted data and display of tabular data in various modules is implemented using DataTables [12]. The interactive outlier-identification step uses Plotly [13]. Scatter-plots, box-plots, and pseudo-time construction in Monocle are done by the ggplot2 package [3,14].

### Back-end variable management

The expression matrix and the metadata sheet are stored separately for each user. The metadata sheet refers to groups, batches, or other properties of the samples in the corresponding expression matrix. All modules share these two types of tables. Other variables shared across all modules include the log-transformed expression matrix, the filtered and normalized expression matrix, the dimensionally reduced matrix, species (human or mouse) and the primary metadata column.

### Batch-effect removal

Batch effect is defined as the unwanted variation introduced in processing or sequencing in potentially different conditions [15]. To remove batch effects, we implement two methods in Granatum: ComBat and Median alignment.

**ComBat:** This method adjusts the batch-effect using empirical Bayes frameworks, and is robust in the presence of outliers or for small sample sizes [16]. It is originally designed for batch-effect removal of microarray gene expression datasets but is commonly used in single-cell RNA-Seq studies [17–19]. It is implemented by the “ComBat” function in the R package “sva” [20].

**Median alignment:** First, this method calculates the median expression of each sample, denoted as *med_i_* for sample *i* Second, it calculates the mean of *med_i_* for each batch, denoted as *batchMean_b_* for batch *b*

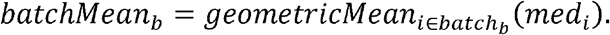

Finally, it multiplies each batch by a factor that pulls the expression levels towards the global geometric mean of the sample medians. When *i* ∈ *batch_b_* and is the number of samples,

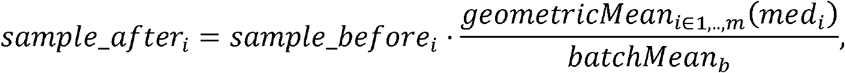

where sample_before_i_ and sample_after_i_ denote the expression levels for all genes within sample *i* before and after batch-effect removal.

### Outlier detection and gene filtering

Z-score threshold is used to automatically detect outliers. The z-score of a cell is calculated by calculating the Euclidean norm of the cell’s vector of expression levels, after scaling all genes to have unit standard deviation and zero mean [21].Over-dispersion gene filtering is done as recommended by Brennecke et al. 2013 [4]. The output of the Monocle package [3] is modified to calculate dispersion and fit a negative binomial model to the result.

### Clustering methods

The following description of clustering algorithms assumes that *n* is the number of genes, is the number of genes, *m* number of samples, and *k* is the number of clusters.

**Non-negative matrix factorization (NMF)**: The log-transformed expression matrix (*n*-by-*m*) is factorized into two non-negative matrices *H*(*n-*by-*m*)and (*k*-by-*m*). The highest-valued *k* entry in each column of determines the membership of each cluster [22,23]. The NMF computation is implemented in the NMF R-package, as reported earlier [22,24].

**K-means:** K-means is done on either the log-transformed expression matrix or the 2-by correlation t-SNE matrix. The algorithm is implemented by the kmeans function in R [25].

**Hierarchical clustering** (Hclust): Hclust is done on either the log-transformed expression matrix or the 2-by-correlation t-SNE matrix. The algorithm is implemented by the hclust function in R [26]. The heatmap with dendrograms is plotted using the heatmap function in R.

### Dimension reduction methods

**Correlation t-SNE:** The method assesses heterogeneity of the data using a two-step process. First, it calculates a distance matrix using the correlation distance. The correlation distance D_i,j_ between sample *i* and sample *j* is defined as

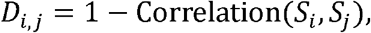

where *S_i_*and *S_j_* are the *i*-th and *j*-th column (sample) of the expression matrix. Next, Rtsne R package [27] uses this distance matrix to reduce the expression matrix to two dimensions.

**PCA:** The Principal Component Analysis algorithm, implemented as “prcomp” function in R, decomposes the original data into linearly uncorrelated variables (components) using orthogonal transformation. The components are then sorted by their variance. The two components with the largest variances (PC1 and PC2) are extracted for visualization [28].

### Elbow-point finding algorithm in clustering

This method is inspired by a similar approach implemented in SCRAT [29]. In the clustering module with automatic determination of the number of clusters, the identification of the optimum number of clusters is done prior to presenting the clustering results. For each number of clusters *k* = 2 to *k* = 10, the percentage of the explained variance (EV) is calculated. To find the elbow-point *k* = *m* where the EV plateaus, a linear elbow function is fit to the *k*-EV data points. This piecewise function consists of a linearly increasing piece from 0 to *m*, and a constant piece from to 10. The algorithm iterates from 1 to 10 and identifies which gives the best coefficient of determination (*R^2^*) of linear regression as the “elbow point”.

### Differential expression analysis

We include four differential expression (DE) algorithms in Granatum: NODES[30], SCDE[31], EdgeR [32], and Limma [33]. Among them, NODES and SCDE are designed for single-cell RNA-Seq specifically. EdgeR and Limma are conventional bulk cell RNA-Seq DE tools that have also been used in single-cell RNA-Seq studies [34,35]. When more than two clusters are present, we perform pairwise DE analysis on all clusters. We use default parameters for all packages. Their versions are: NODES (0.0.0.9010), SCDE (1.99.2), EdgeR (3.18.1) and Limma (3.32.2)

### Gene-set enrichment analysis

The *fgsea* R-package implements the Gene Set Enrichment Analysis (GSEA) algorithm with optimizations for speedup [36,37]. GSEA calculates an enrichment score, which quantifies the relevance of a gene set (for example, a KEGG pathway or a GO term) to a particular group of selected genes (e.g., DE genes called by a method). The p-value is calculated for each gene set according to the empirical distribution, followed by Benjamini–Hochberg multiple hypothesis tests [38].

### Pseudo-time construction

We use Monocle (version 2.2.0) in our pseudo-time construction step. When building the CellDataSet required for monocle’s input, we set the *expressionFamily to negbinomial.size()*. We use *reduceDimension* function to reduce the dimensionality by setting *max_components* to 2.

## Results

### Overview of Granatum

Granatum is by far the most comprehensive graphic-user-interface (GUI) based scRNA-Seq analysis pipeline with no requirement of programming knowledge (Table 1). It allows both direct web-based analysis (accessible through either desktop computers or mobile devices), as well as local deployment (as detailed in the front-page of http://garmiregroup.org/granatum/app). The project is fully open source, and its source code can be found at http://garmiregroup.org/granatum/code.

**Table 1.** Comparison of existing single-cell analysis pipelines.

We have systematically compared Granatum with 12 other existing tools to demonstrate its versatile functions (Table 1). Popular packages such as SCDE / PAGODA and Flotilla are developed for programmers and require expertise in a particular programming language. In contrast, Granatum with its easy-to-navigate graphical interface requires no programming specialty. The current version of Granatum neatly presents nine modules, arranged as steps and ordered by their dependencyIt starts with one or more expression matrices and corresponding sample metadata sheet(s), followed by data merging, batch-effect removal, outlier removal, normalization, gene filtering, clustering, differential expression, protein-protein network, and pseudo-time construction.

Besides the features above, a number of enhanced functionalities make Granatum more flexible than other freely available tools (Table 1). (1) Unlike tools such as SCRAT (https://zhiji.shinyapps.io/scrat/), ASAP [39] and Sake (http://sake.mhammell.tools/), it is the only GUI pipeline that supports multiple dataset submission as well as batch effect removal. (2) Each step can be reset for re-analysis. (3) Certain steps (eg. batch-effect removal, outlier removal, and gene filtering) can be bypassed without affecting the completion of the workflow. (4) Subsets of the data can be selected for customized analysis. (5) Outlier samples can be identified either automatically by a pre-set threshold or by manually clicking/lassoing the samples the PCA plot or the correlation t-SNE plot. (6) Multiple cores can be utilized in the differential expression module for speed-up. (7) Both GSEA and network analysis can be performed for the differentially expressed genes in all pairs of subgroups, following clustering analysis. (8) Pseudo-time construction is included, giving insights into relationships between the cells.

### Testing of the software

In this report, we mainly use a previously published data set as an example [18]. This renal carcinoma dataset contains three groups of cells: patient-derived xenografts (PDX) primary, PDX metastatic cells, and patient metastatic cells [18]. We abbreviate this dataset as the K-dataset.

To estimate the total running time of Granatum (with default parameters) at different sizes of datasets, we first simulate expression matrices with 200, 400, 800, or 1600 cells using the Splatter package, based on the parameters estimated from the K-dataset [40]. Additionally, we also use down-sample approach (200, 400, 800, 1600, 3200 and 6000 cells) on a dataset (P-dataset) provided by 10x Genomics, which has 6,000 peripheral blood mononuclear cells (PBMCs) (https://support.10xgenomics.com/single-cell-gene-expression/datasets/1.1.0/pbmc6k). The running time scales linearly with the number of cells, regardless of platform (Suppl. Figure 1). The most time-consuming step is Monocle based pseudo-time construction, which takes about 80% of all computing time.

**Figure 1.**
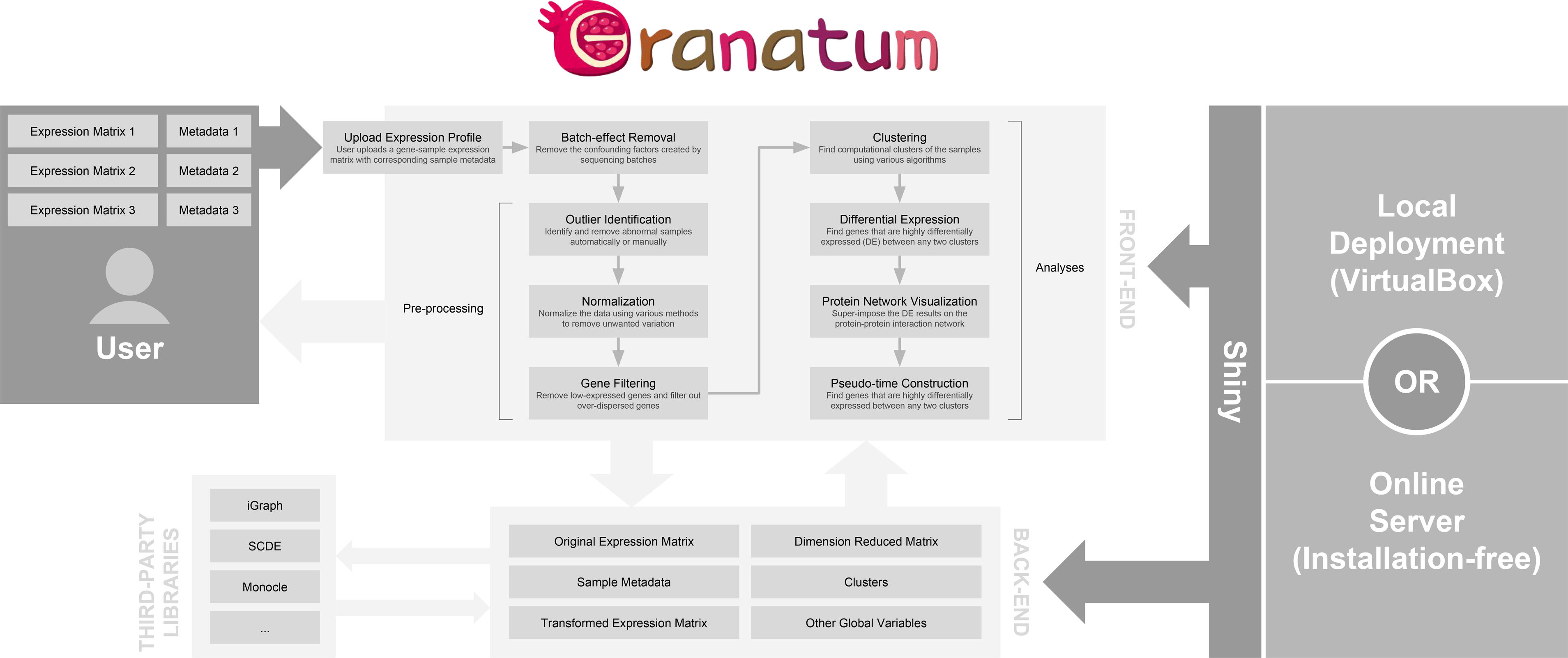
Granatum workflow. Granatum is built with the Shiny framework, which integrates the front-end with the back-end. A public server has been provided for easy access, and local deployment is also possible. The user uploads one or more expression matrices with corresponding metadata for samples. The back-end stores data separately for each individual user, and invokes third-party libraries on demand.

In the following sections, we use K-dataset to elaborate the details of each step in Granatum in chronological order.

### Upload data

Granatum accepts one or more expression matrices as input. Each expression matrix may be accompanied by a metadata sheet. A metadata sheet is a table describing the groups, batches, or other properties of the samples in the corresponding expression matrix. Users may upload multiple matrices sequentially. Currently, Granatum accepts either human or mouse species, for downstream functional analysis. After uploading the input files, users can preview the matrix and metadata tables to validate that the dataset is uploaded correctly.

### Batch-effect removal

Samples obtained in batches can create unwanted technical variation, which confounds the biological variation [15]. It is therefore important to remove the expression level difference due to batches. Granatum provides a batch-effect removal step where two methods are included, namely ComBat [16] and median alignment. If multiple datasets are uploaded, by default, each dataset is assumed to be one batch. Alternatively, if the batch numbers are indicated in the sample metadata sheet, the user may select the column in which the batch numbers are stored. For datasets with a large number of cells, the box-plot shows a random selection of 96 sub-samples for the visualization purpose and can be re-sampled freely.

To show that median alignment can effectively remove the batches, we randomly select half of the cells in K-dataset and multiply the expression levels by 3, thus creating two artificial batches 1 and 2. The PCA plot shows that due to the batch-effect, cells of the same type are separated by batch (the two colors) (Figure 2A). After performing median alignment, the batch effect is minimized, and cells from the same type but in two colors (batches) are now intermingled (Figure 2B).

**Figure 2.**
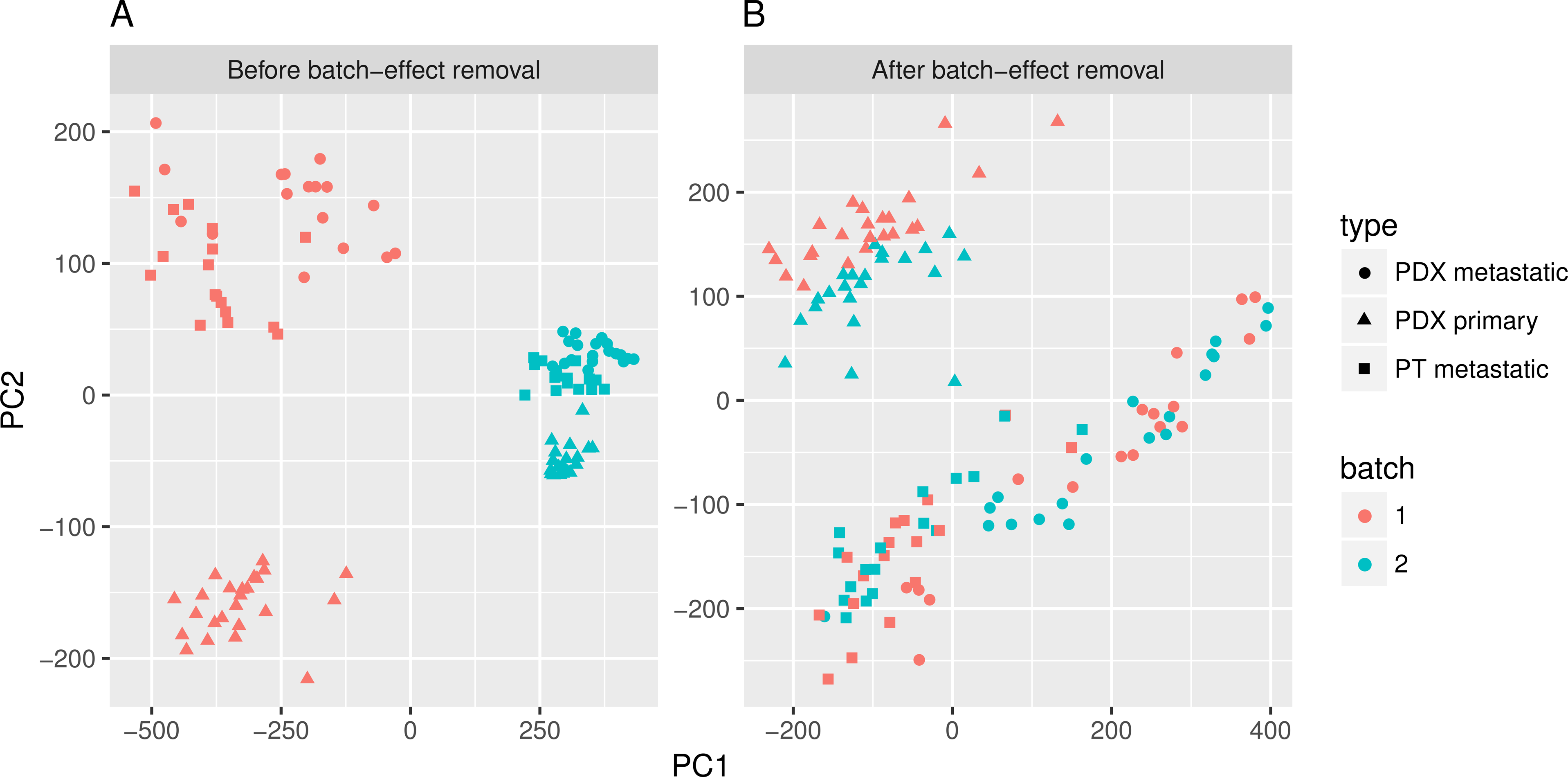
The batch-effect removal. The PCA plots show the before/after median alignment comparison. The colors indicate the two batches 1 and 2, and the shapes indicate the three cell types reported from the original data. (A) Before and (B) After batch-effect removal.

### Outlier identification

Computationally abnormal samples pose serious problems for many downstream analysis procedures. Thus, it is crucial to identify and remove them in the early stage. Granatum’s outlier identification step features PCA and t-SNE [41] plots, two connected interactive scatter-plots that have different computational characteristics. A PCA plot illustrates the Euclidean distance between the samples, and a correlation t-SNE plot shows the associative distances between the samples. Granatum generates these two plots using top genes (default 500). Using the Plotly library [13], these plots are highly interactive. It is an example of thoughtful tool design that empowers users to explore the data. Outliers can be identified automatically by using a z-score threshold or setting a fixed number of outliers. In addition, each sample can be selected or de-selected, by clicking, boxing or drawing a lasso on its corresponding points.

The original K-dataset has one sample with abnormally low expression level. This potential outlier sample can affect downstream analyses. Using Granatum, users can easily spot such outliers in the PCA plot or in the correlation t-SNE plot (Figure 3A and B). After removal of the outliers, the top-gene based PCA and correlation t-SNE plots are more balanced (Figure 3C and D).

**Figure 3.**
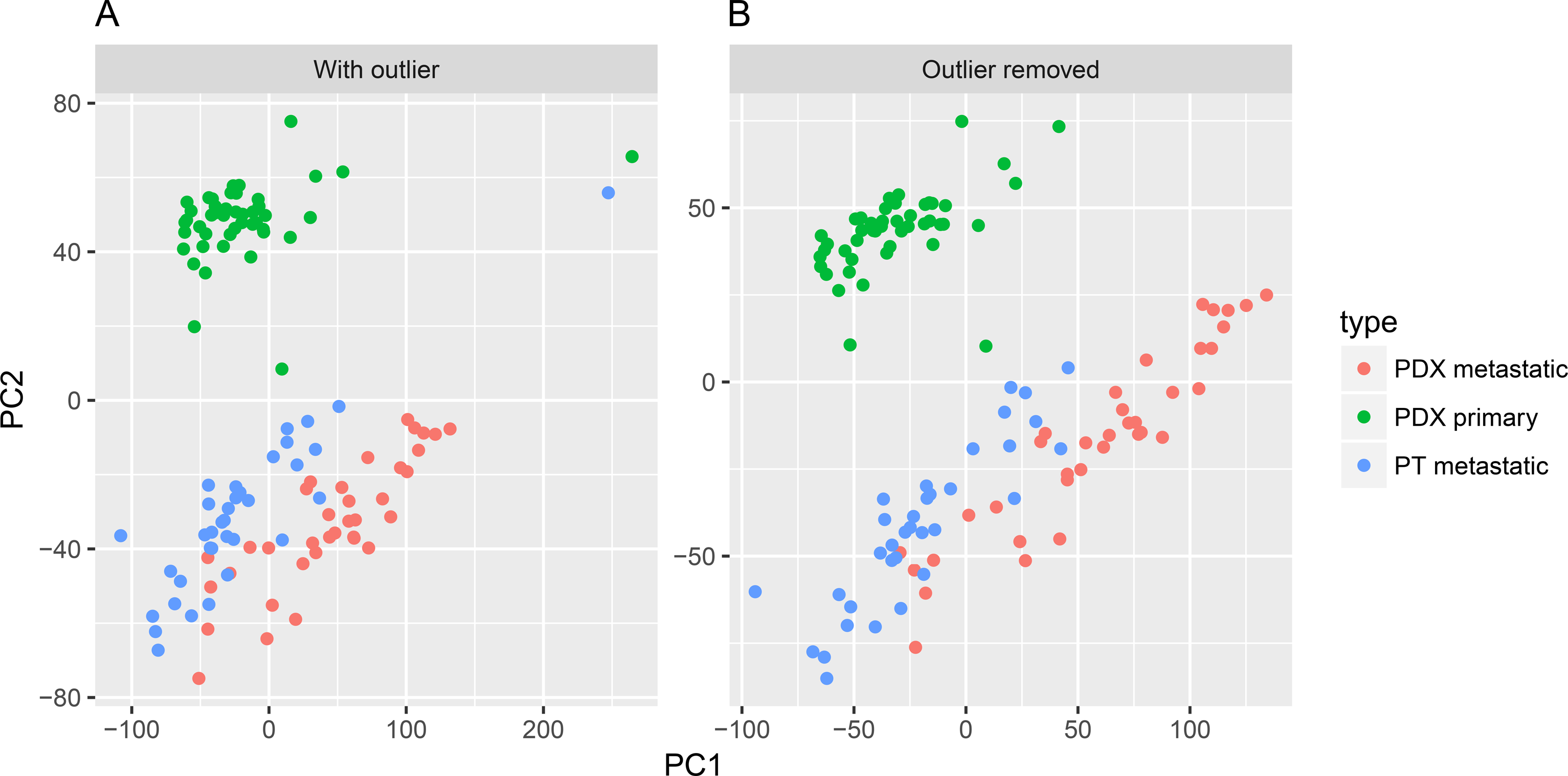
The outlier removal using PCA plot. (A) Before outlier removal. (B) After outlier removal.

### Normalization

Normalization is essential to most scRNA-Seq data before the downstream functional analyses (except those with the UMI counts). Granatum includes four commonly used normalization algorithms: quantile normalization, geometric mean normalization, size-factor normalization [42,43], and Voom [44]. A post-normalization box-plot helps illustrate the normalization effect to the median, mean, and extreme values across samples.

The box-plots allow observation of various degrees of stabilization (Figure 4). The original dataset has high levels variations among samples (Figure 4A). Quantile normalization unifies the expression distribution of all samples, thus renders the box-plots identical (Figure 4B). Mean alignment tries to unify all means of the samples by multiplying the expression levels in each sample by a factor, thus visually all means (the red dots) are the same (Figure 4C). Size-factor and Voom normalization use more sophisticated procedures to normalize the data, but the variation of distribution across samples is evidently reduced (Figure 4D and E). According to our experience and others [45,46], quantile normalization is recommended.

**Figure 4.**
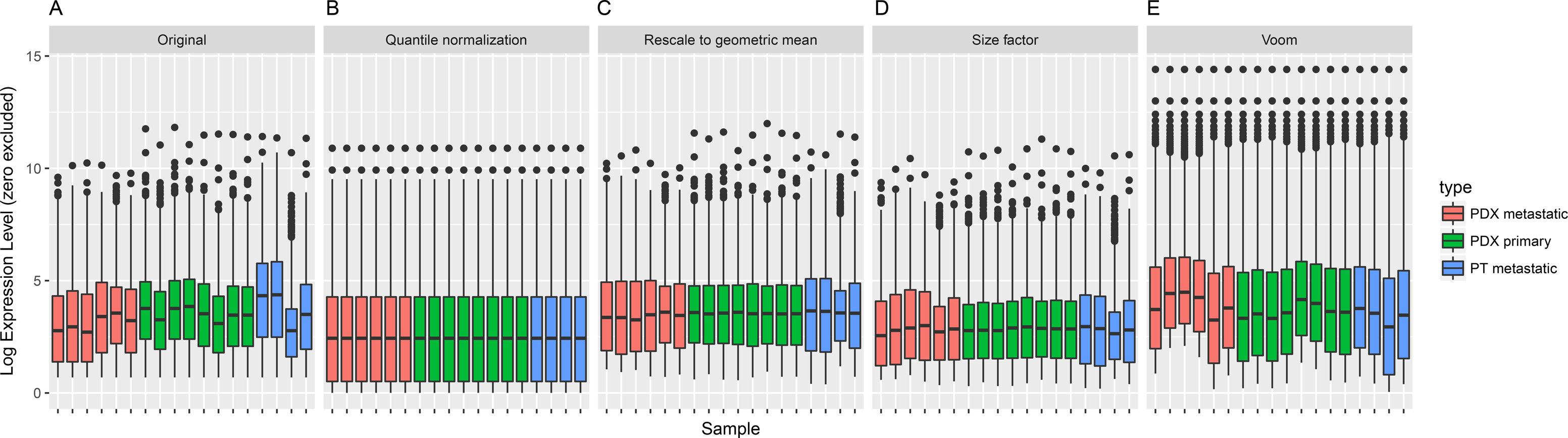
Box-plot comparison of normalization methods. The cells size is down-sampled to representatively show the general effect of each method. The colors indicate the three cell types reported from the original data. (A) The original (no normalization) (B) Quantile normalization (C) Geometrical mean normalization (D) Size-factor normalization (E) Voom normalization.

### Gene filtering

Due to high noise levels in scRNA-Seq, Brennecke et al. [4] recommended removing lowly expressed genes as well as lowly-dispersed genes. To this end, Granatum includes a step to remove these genes. Both the average expression-level threshold and the dispersion threshold can be adjusted interactively. Granatum displays the threshold selection sliders and the number-of-genes statistics message to enhance integration with the other components. On the mean-dispersion plot, a point represents a gene, whose x-coordinate is the log transformed mean of the expression levels of that gene, and the y-coordinate is the dispersion factor calculated from a negative binomial model. The plot highlights the preserved genes as black and the filtered genes as gray (Suppl. Figure 2).

### Clustering

Clustering is a routine heuristic analysis for scRNA-Seq data. Granatum selects five commonly used algorithms: non-negative matrix factorization [22], k-means, k-means combined with correlation t-SNE, hierarchical clustering (hclust), and hclust combined with correlation t-SNE. The number of clusters can either be set manually, or automatically using an elbow-point finding algorithm. For the latter automatic approach, the algorithm will cluster samples with the number of clusters (*k*) ranging from 2 to 10, and determine the best number as the elbow-point *k*. the starting point of the plateau for explained variance (EV). If hclust is selected, a pop-up window shows a heatmap with hierarchical grouping and dendrograms.

Next, the two unsupervised PCA and correlation t-SNE plots superimpose the resulting cluster labels on the samples (Suppl. Figure 3). Users can also chose to use their pre-defined labels provided in the sample metadata. By comparing the two sets of labels, one can check the agreement between the prior metadata labels and the computed clusters. We perform the K-means clustering (*k* = 2) on the correlation t-SNE plot, using K-dataset. The generated clusters perfectly correspond to the original cell type labels in this case.

### Differential expression

After the clustering step, Granatum allows DE analysis on genes between any two clusters. It currently includes four commonly used differential expression methods, namely NODES [30], SCDE [31], Limma [33] and edgeR [32]. The DE analysis is performed in a pair-wise fashion when more than two clusters are present. To shorten the computation time, the number of cores for parallelization on multi-core machines can be selected. When the DE computation is complete, the results are shown in a table with DE genes sorted by their Z-scores, along with the coefficients. As another feature to empower the users, the gene symbols are linked to their corresponding GeneCards pages (www.genecards.org) [47]. The “Download CSV table” button allows saving the DE results as a CSV file.

Next, Gene Set Enrichment Analysis (GSEA) with either KEGG pathways or Gene Ontology (GO) terms [37,48–50] can be performed, to investigate the biological functions of these DE genes. The results are plot in an intuitive bubble-plot (Figure 5D). In this plot, the y-axis represents the enrichment score of the gene sets, x-axis shows gene set names, and the size of the bubble indicates the number of genes in that gene set.

**Figure 5.**
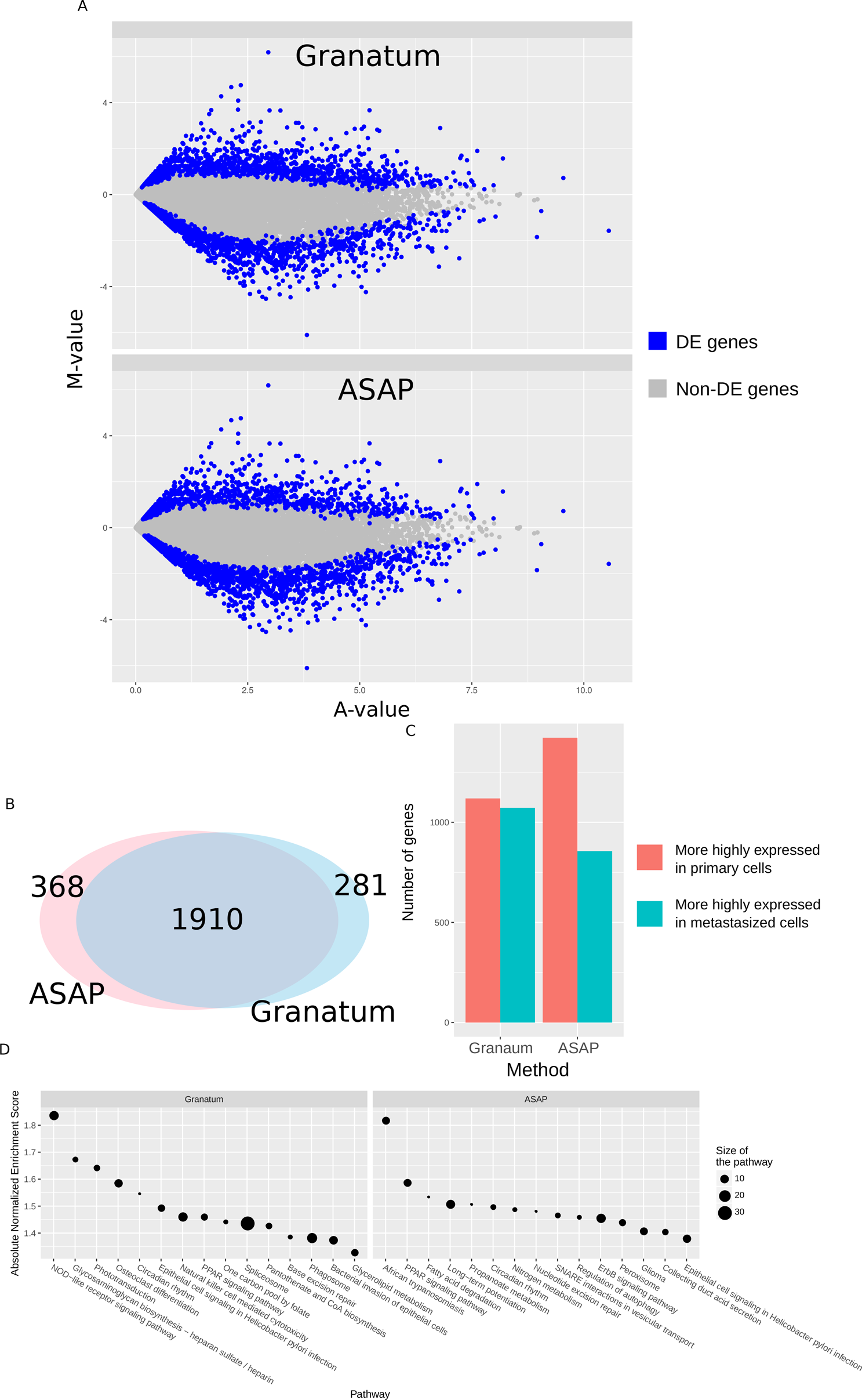
Comparison of DE genes identified by Granatum or ASAP pipeline. (A) MA-plot. Blue color labels DE genes, and gray dots are non-DE genes. (B) Venn diagram showing the number of DE genes identified by both methods, as well as those uniquely identified by either pipeline. (C) Bar chart comparing the number of genes up regulated in primary cells (red) or metastasized cells (green). (D) Bubble-plots of KEGG pathway GSEA results for the DE genes identified by either pipeline. The y-axis represents the enrichment score of the gene sets, x-axis shows gene set names, and the size of the bubble indicates the number of genes in that gene set.

### Comparison with other Graphical web tools of scRNA-Seq

To evaluate the differences between Granatum and a similar graphical scRNA-Seq pipeline ASAP [39], we compare the DE genes (primary vs. metastasized patient) in K-dataset obtained by both pipelines (Figure 5). While Granatum uses quantile normalization, ASAP uses Voom normalization as default method. We use SCDE as it is the common DE method for both pipelines.

Both pipelines agree on most DE genes called (Figure 5A), but each identifies a small number of unique DE genes (Figure 5B). In Granatum, the number of up or down regulated DE genes detected by Granatum are closer. Whereas in ASAP, a lot more genes are higher regulated in the primary cells, compared to those in metastasized cells (Figure 5C). Further, KEGG pathway based GSEA analysis on the DE genes shows that Granatum identified more significantly (Enrichment Score > 1.5) enriched pathways than ASAP (Figure 5C). The top pathway enriched in Granatum’s DE genes is the NOD-like receptor-signaling pathway, corresponding to its known association with immunity and inflammation [51]. In ASAP “African trypanosomiasis” is the top pathway, which describes the molecular events when parasite Trypanosoma brucei pass through the blood-brain barrier and cause neurological damage by inducing cytokines. Despite the differences, some signaling pathways are identified by both pipelines with known associations with tumorigenesis, such as PPAR signaling pathway [52] and Epithelial cell signaling pathway [53].

### Granatum-specific Steps: Protein network visualization and Pseudo-time construction

Unlike ASAP, SAKE and SCRAT, Granatum implements a Protein-protein interaction (PPI) network to visualize the connections between the DE genes (Figure 6A). By default, up to 200 genes are displayed in PPI network. We use visNetwork to enable the interactive display of the graph [11], so that users can freely rearrange the graph by dragging the nodes to the desired location. Uses can also reconfigure the layout to achieve good visualization via an elastic-spring physics simulation. Nodes are colored according to their regulation direction and the amount of change (quantified using Z-score), where red indicates up-regulation and blue indicates down-regulation. As an example, Figure 6A shows the PPI network result from PDX primary to metastatic cells in the K dataset. A large, closely connected module exists in PPI network, which contains many heat shock protein genes including down-regulated HSP90AB1, HSPA6, HSPA7, HSPA8, HSPA1A, HSPA1B and HSPA4L as well as up-regulated HSP90AA1 and HSPH1 in metastasized cells. Heat shock genes have been long recognized as a stress response genes [54], and inhibiting heat shock protein genes can control metastasis in various types of cancers [55,56].

**Figure 6.**
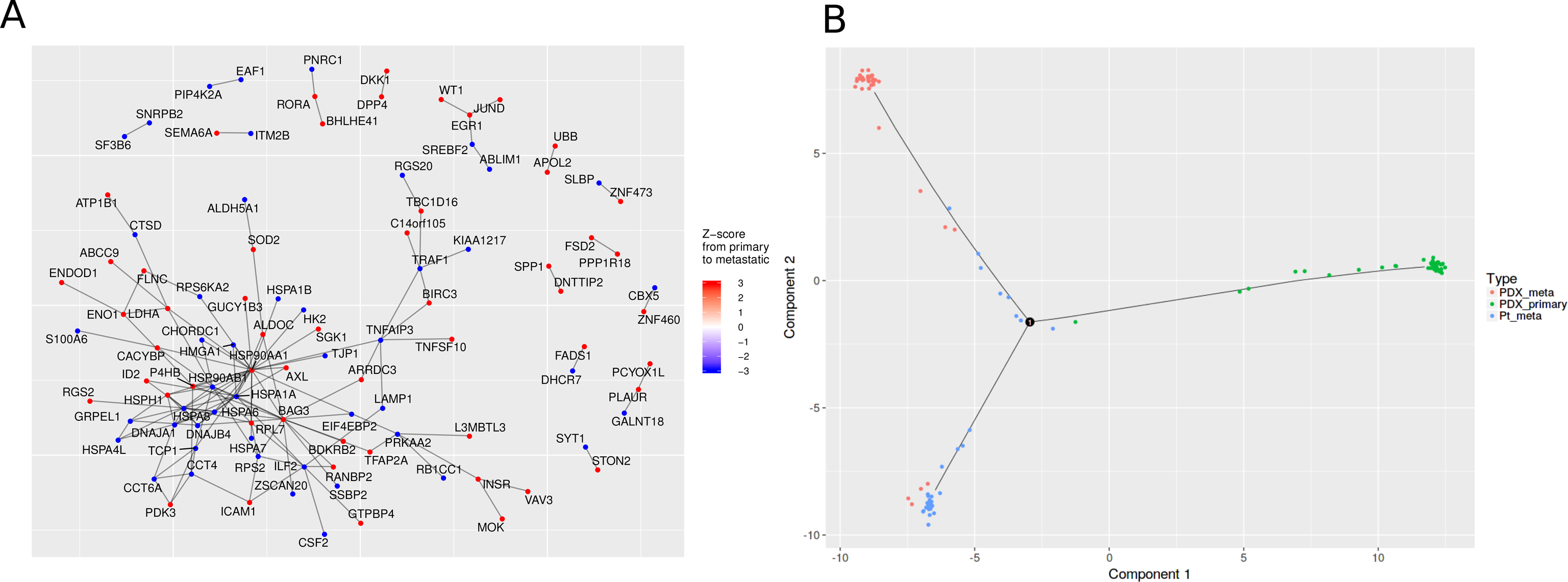
The Protein-protein interaction network and Pseudo-time construction steps. (A) The PPI network derived from the DE results between PDX primary and metastasized cells in the K-dataset. The color on each node (gene) indicates its Z-score in the differential expression test. Red and blue colors indicate up- and down-regulation in metastasized cells, respectively. (B) The Pseudo-time construction step. Monocle algorithm is customized to visualize the paths among individual cells. Sample labels from the metadata are shown as different colors in the plot.

Lastly, Granatum has included the Monocle algorithm[3], a widely-used method to reconstruct a pseudo-timeline for the samples (Figure 6B). Monocle uses the Reversed Graph Embedding algorithm to learn the structure of the data, as well as the Principal Graph algorithm to find the timelines and branching points of the samples. The user may map any pre-defined labels provided in the metadata sheet onto the scatter-plot. In the K-dataset, the three (PDX primary, PDX metastasized, and patient metastasized) types of cancer cells are mostly distinct (Figure 6B). However, small portions of cells from each type appear to be on intermediate trajectory.

## Discussion

The field of scRNA-Seq is fast-evolving both in terms of the development of instrumentation and the innovation of computational methods. However, it becomes exceedingly hard for a wet-lab researcher without formal bioinformatics training to catch up with the latest iterations of algorithms [5]. This barrier forces many researchers to resort to sending their generated data to third-party bioinformaticians before they are able to visualize the data themselves. This segregation often prolongs the research cycle time, as it often takes significant effort to maintain effective communications between wet-lab researchers and bioinformaticians. In addition, issues with the experimentations do not get the chance to be spotted early enough to avoid significance loss of time and cost in the projects. It is thus attractive to have a non-programming graphical application that includes state-of-the-art algorithms as routine procedures, in the hands of the bench-scientist who generate the scRNA-Seq data.

Granatum is our attempt to fill this void. It is to our knowledge the most comprehensive solution that aims to cover the entire scRNA-Seq workflow with an intuitive graphical user interface. Throughout the development process, our priority has been to make sure that it is fully accessible to researchers with no programming experiments. We have strived to achieve this, by making the plots and tables self-explanatory, interactive and visually pleasant. We have sought inputs from our single-cell bench-side collaborators to ensure that the terminologies are easy to understand by them. We also supplement Granatum with a manual and online video that guide the users through the entire workflow, using example datasets. We also seek feedback from community via Github pull-requests, emails discussions and user survey.

Currently, Granatum targets bench scientists who have their expression matrices and metadata sheets ready. However, we are developing the next version of Granatum, which will handle the entire scRNA-Seq data processing and analysis pipeline including FASTQ quality control, alignment, and expression quantification. Another caveat is the lacking of benchmark dataset in single-cell analysis field currently, where the different computational packages can be evaluated unbiasedly. We thus resort to empirical comparisons on packages between Granatum and ASAP. In the future, we will enrich Granatum with capacities to analyze and integrate other types of genomics data in single cells, such as exome-seq and methylation data. We will closely update Granatum to keep up with the newest development in the scRNA-Seq bioinformatics field. We welcome third-party developers to download the source-code and modify Granatum, and will continuous integrate and innovate this tool as the go-to place for single-cell bench scientists.

## Conclusions

We have developed a graphical web application called Granatum, which enables bench researchers with no programming expertise to analyze state-of-the-art scRNA-Seq data. This tool offers many interactive features to allow routine computational procedures with a great amount of flexibility. We expect that this platform will empower the bench-side researchers with more independence in the fast-evolving single cell genomics field.

## Declarations

### Ethics approval and consent to participate

Not Applicable.

### Consent for publication

Not Applicable.

### Availability of data and material

All datasets used in the comparisons are reported by previous studies. The K-dataset has the NCBI Gene Expression Omnibus (GEO) accession number GSE73122. The 6,000 cells PBMCs dataset is retried from 10x Genomics website https://support.10xgenomics.com/single-cell-gene-expression/datasets/1.1.0/pbmc6k.

Granatum can be visited at: http://garmiregroup.org/granatum/app

Granatum source-code can be found at: http://garmiregroup.org/granatum/code

A demonstration video can be found at: http://garmiregroup.org/granatum/video

## Supplementary Figures

**Suppl. Figure 1: Granatum total running time with various numbers of cells.** Datasets with various sizes from two single-cell platforms (Fluidigm C1 and 10x Genomics) are used. To generate expression data up to 6000 cells, the Fluidigm C1 datasets are simulated using Splatter, with parameters estimated from the K-dataset (118 cells). The 10x Genomics datasets are down-sampled from the original 6000-cell PBMC dataset. The x-axis represents the size of the dataset, and the y-axis represents the total running time (in minutes) of Granatum. Monocle based pseudo-time construction step takes about 80% of total running time.

**Suppl. Figure 2: The Gene filtering step.** The y-axis of the scatter-plot is the empirical dispersion, estimated by a negative binomial model. The x-axis is the log mean expression of each gene. The red line is the fit of a negative binomial model onto the data. Black points represent gene to be kept and gray points are filtered genes.

**Suppl. Figure 3: The Clustering step.** (A) PCA and (B) Correlation t-SNE plots of single cells (dots) are shown, with colors indicating the cell types reported in the original dataset and cluster number (1, 2) super-imposed on the cells.

## Authors’ contributions

LXG envisioned the project. XZ developed the majority of the pipeline. TW and AT assisted in developing the pipeline. TW documented the user manual and performed packaging. XZ, TW and LXG wrote the manuscript. All authors have read, revised, and approved the final manuscript.

## Acknowledgements

We thank Drs. Michael Ortega and Paula Benny for providing valuable feedback during testing the tool. We also thank other group members in Garmire group for suggestions in the tool development.

## List of abbreviations

scRNA-Seq: Single-cell high-throughput RNA sequencing
DE: differential expression
GSEA: Gene-set enrichment analysis
KEGG: Kyoto Encyclopedia of Genes and Genomes
GO: Gene ontology
PCA: Principal component analysis
SNE: t-Distributed Stochastic Neighbor Embedding
NMF: Non-negative matrix factorization
Hclust: Hierarchical clustering
PPI: Protein-protein interaction

## References

1. Patel AP Tirosh I Trombetta JJ Shalek AK Gillespie SM Wakimoto H et al. Single-cell RNA-seq highlights intratumoral heterogeneity in primary glioblastoma. Science (80-.). 2014;344:1396–401.

2. Lewis BP Burge CB Bartel DP. Conserved seed pairing often flanked by adenosines indicates that thousands of human genes are microRNA targets. Cell. Elsevier; 2005;120:15–20.

3. Trapnell C Cacchiarelli D Grimsby J Pokharel P Li S Morse M et al. The dynamics and regulators of cell fate decisions are revealed by pseudotemporal ordering of single cells. Nat. Biotechnol. Nature Research; 2014;32:381–6.

4. Brennecke P Anders S Kim JK Kołodziejczyk AA Zhang X Proserpio V et al. Accounting for technical noise in single-cell RNA-seq experiments. Nat. Methods. Nature Publishing Group; 2013;

5. Poirion OB Zhu X Ching T Garmire L. Single-Cell Transcriptomics Bioinformatics and Computational Challenges. Front. Genet. 2016. p. 163.

6. Team RC. R: A language and environment for statistical computing. R Foundation for Statistical Computing Vienna, Austria. 2015, URL http://www.R-project.org. 2016;

7. McCarthy DJ Campbell KR Lun ATL Wills QF. scater: pre-processing, quality control normalisation and visualisation of single-cell RNA-seq data in R. bioRxiv [Internet]. Cold Spring Harbor Labs Journals; 2016; Available from: http://biorxiv.org/content/early/2016/08/15/069633

8. Ihaka R Gentleman R. R: a language for data analysis and graphics. J. Comput. Graph. Stat. Taylor & Francis; 1996;5:2 99–314.

9. RStudio, Inc. Easy web applications in R. 2013.

10. Attali D. shinyjs: Easily Improve the User Experience of Your Shiny Apps in Seconds [Internet]. 2016. Available from: https://cran.r-project.org/package=shinyjs

11. Almende B.V., Thieurmel B. visNetwork: Network Visualization using “vis.js” Library [Internet]. 2016. Available from: https://cran.r-project.org/package=visNetwork

12. Xie Y. DT: A Wrapper of the JavaScript Library “DataTables” [Internet]. 2016. Available from: https://cran.r-project.org/package=DT

13. Sievert C Parmer C Hocking T Chamberlain S Ram K Corvellec M et al. plotly: Create Interactive Web Graphics via “plotly.js” [Internet]. 2016. Available from: https://cran.rproject.org/package=plotly

14. Wickham H. ggplot2: Elegant Graphics for Data Analysis [Internet]. Springer-Verlag New York; 2009. Available from: http://ggplot2.org

15. Hicks SC Teng M Irizarry RA. On the widespread and critical impact of systematic bias and batch effects in single-cell RNA-Seq data. bioRxiv. Cold Spring Harbor Labs Journals; 2015;25528.

16. Johnson WE Li C Rabinovic A. Adjusting batch effects in microarray expression data using empirical Bayes methods. Biostatistics. Biometrika Trust; 2007;8:118–27.

17. Kim K-T, Lee HW Lee H-O, Kim SC Seo YJ Chung W et al. Single-cell mRNA sequencing identifies subclonal heterogeneity in anti-cancer drug responses of lung adenocarcinoma cells. Genome Biol. 2015;16:127.

18. Kim K-T, Lee HW Lee H-O, Song HJ Shin S Kim H et al. Application of single-cell RNA sequencing in optimizing a combinatorial therapeutic strategy in metastatic renal cell carcinoma. Genome Biol. BioMed Central; 2016;17:80.

19. Petropoulos S Edsgärd D Reinius B Deng Q Panula SP Codeluppi S et al. Single-Cell RNA-Seq Reveals Lineage and X Chromosome Dynamics in Human Preimplantation Embryos. Cell. Elsevier; 2016;

20. Leek JT Storey JD. Capturing heterogeneity in gene expression studies by surrogate variable analysis. PLoS Genet. Public Library of Science; 2007;3:e161.

21. Iglewicz B Hoaglin DC. How to detect and handle outliers. Asq Press; 1993.

22. Zhu X Ching T Pan X Weissman S Garmire L. Detecting heterogeneity in single-cell RNA-Seq data by non-negative matrix factorization. PeerJ Prepr. PeerJ Inc. San Francisco USA; 2016;4:e1839v1.

23. Brunet J-P, Tamayo P Golub TR Mesirov JP. Metagenes and molecular pattern discovery using matrix factorization. Proc. Natl. Acad. Sci. 2004;101:4164–9.

24. Gaujoux R Seoighe C. Algorithms and framework for nonnegative matrix factorization (NMF). 2010.

25. Lloyd S. Least squares quantization in PCM. IEEE Trans. Inf. theory. IEEE; 1982;28:129–37.

26. Murtagh F Contreras P. Methods of hierarchical clustering. arXiv Prepr. arXiv1105.0121. 2011;

27. Krijthe J. Rtsne: T-Distributed Stochastic Neighbor Embedding using Barnes-Hut Implementation. R Packag. version 0.10, URL http://CRAN.R-project.org/package=Rtsne. 2015;

28. Pearson K. LIII. On lines and planes of closest fit to systems of points in space. London, Edinburgh, Dublin Philos. Mag. J. Sci. Taylor & Francis; 1901;2:559–72.

29. Ji Z Zhou W Ji H. Single-cell regulome data analysis by SCRAT. Bioinformatics. Oxford University Press; 2017;btx315.

30. Sengupta D Rayan NA Lim M Lim B Prabhakar S. Fast, scalable and accurate differential expression analysis for single cells. bioRxiv. Cold Spring Harbor Labs Journals; 2016;49734.

31. Kharchenko P V Silberstein L Scadden DT. Bayesian approach to single-cell differential expression analysis. Nat. Methods. Nature Publishing Group; 2014;11:740–2.

32. Robinson MD McCarthy DJ Smyth GK. edgeR: a Bioconductor package for differential expression analysis of digital gene expression data. Bioinformatics. Oxford Univ Press; 2010;26:139–40.

33. Ritchie ME Phipson B Wu D Hu Y Law CW, Shi W et al. limma powers differential expression analyses for RNA-sequencing and microarray studies. Nucleic Acids Res. Oxford University Press; 2015;43:e47–e47.

34. Fan X Zhang X Wu X Guo H Hu Y Tang F et al. Single-cell RNA-seq transcriptome analysis of linear and circular RNAs in mouse preimplantation embryos. Genome Biol. BioMed Central; 2015;16:148.

35. Tasic B Menon V Nguyen TN Kim TK Jarsky T Yao Z et al. Adult mouse cortical cell taxonomy by single cell transcriptomics. Nat. Neurosci. NIH Public Access; 2016;19:335.

36. Sergushichev A. An algorithm for fast preranked gene set enrichment analysis using cumulative statistic calculation. bioRxiv [Internet]. Cold Spring Harbor Labs Journals; 2016; Available from: http://biorxiv.org/content/early/2016/06/20/060012

37. Subramanian A Tamayo P Mootha VK Mukherjee S Ebert BL Gillette MA et al. Gene set enrichment analysis: a knowledge-based approach for interpreting genome-wide expression profiles. Proc. Natl. Acad. Sci. National Acad Sciences; 2005;102:15545–50.

38. Benjamini Y Hochberg Y. Controlling the false discovery rate: a practical and powerful approach to multiple testing. J. R. Stat. Soc. Ser. B. JSTOR; 1995;289–300.

39. Gardeux V David F Shajkofci A Schwalie PC Deplancke B. ASAP: a Web-based platform for the analysis and inter-active visualization of single-cell RNA-seq data. bioRxiv. Cold Spring Harbor Labs Journals; 2016;96222.

40. Zappia L Phipson B Oshlack A. Splatter: Simulation Of Single-Cell RNA Sequencing Data. bioRxiv. Cold Spring Harbor Labs Journals; 2017;133173.

41. Maaten L van der Hinton G. Visualizing data using t-SNE. J. Mach. Learn. Res. 2008;9:2579–605.

42. Bolstad BM Irizarry RA Åstrand M Speed TP. A comparison of normalization methods for high density oligonucleotide array data based on variance and bias. Bioinformatics. Oxford Univ Press; 2003;19:185–93.

43. Love MI Huber W Anders S. Moderated estimation of fold change and dispersion for RNA-Seq data with DESeq2. bioRxiv. Cold Spring Harbor Labs Journals; 2014;

44. Law CW Chen Y Shi W Smyth GK. Voom: precision weights unlock linear model analysis tools for RNA-seq read counts. Genome Biol. BioMed Central; 2014;15:R29.

45. Xue Z Huang K Cai C Cai L Jiang C Feng Y et al. Genetic programs in human and mouse early embryos revealed by single-cell RNA sequencing. Nature. NIH Public Access; 2013;500:593.

46. Hansen KD Irizarry RA Wu Z. Removing technical variability in RNA-seq data using conditional quantile normalization. Biostatistics. Oxford University Press; 2012;13:204–16.

47. Rebhan M Chalifa-Caspi V Prilusky J Lancet D. GeneCards: integrating information about genes proteins and diseases. Trends Genet. Elsevier Current Trends; 1997;13:163.

48. Kanehisa M Furumichi M Tanabe M Sato Y Morishima K. KEGG: new perspectives on genomes pathways, diseases and drugs. Nucleic Acids Res. Oxford Univ Press; 2017;45:D353–D361.

49. Consortium GO others. Gene ontology consortium: going forward. Nucleic Acids Res. Oxford Univ Press; 2015;43:D1049–D1056.

50. Ashburner M Ball CA Blake JA Botstein D Butler H Cherry JM et al. Gene Ontology: tool for the unification of biology. Nat. Genet. Nature Publishing Group; 2000;25:25–9.

51. Fritz JH Ferrero RL Philpott DJ Girardin SE. Nod-like proteins in immunity inflammation and disease. Nat. Immunol. Nature Publishing Group; 2006;7:1250–7.

52. Belfiore A Genua M Malaguarnera R. PPAR-agonists and their effects on IGF-I receptor signaling: implications for cancer. PPAR Res. Hindawi Publishing Corporation; 2009;2009.

53. Watkins DN Berman DM Burkholder SG Wang B Beachy PA Baylin SB. Hedgehog signalling within airway epithelial progenitors and in small-cell lung cancer. Nature. Nature Publishing Group; 2003;422:313–7.

54. Santoro MG. Heat shock factors and the control of the stress response. Biochem. Pharmacol. Elsevier; 2000;59:55–63.

55. Tamura Y Peng P Liu K Daou M Srivastava PK. Immunotherapy of tumors with autologous tumor-derived heat shock protein preparations. Science (80-.). American Association for the Advancement of Science; 1997;278:117–20.

56. Eccles SA Massey A Raynaud FI Sharp SY Box G Valenti M et al. NVP-AUY922: a novel heat shock protein 90 inhibitor active against xenograft tumor growth angiogenesis, and metastasis. Cancer Res. AACR; 2008;68: 2850–60.

